# Cellular-level control on global ocean deoxygenation driven by phytoplankton ecophysiology

**DOI:** 10.1101/2025.10.08.681239

**Authors:** Shlomit Sharoni, Keisuke Inomura, Stephanie Dutkiewicz, Oliver Jahn, Gregory L Britten, Michael J Follows

## Abstract

Phytoplankton elemental composition shapes the distribution of dissolved nutrient concentrations and thus plays a key role in ocean biogeochemistry. While the carbon-to-nitrogen-to-phosphorus ratio (C:N:P) in marine phytoplankton has been extensively studied, the hydrogen (H) and oxygen (O) content have received less attention despite their critical role in determining dissolved oxygen (O_2_) consumption rates in the ocean. Here, we estimated the elemental composition of marine phytoplankton, including the H and O content, from first principles, using a cellular allocation model embedded in a global ocean model. We estimated that an average phytoplankton cell has a chemical formula of C_107_H_190_N_16_O_53_P, with an O_2_ demand of 149 mol O_2_/mol P and respiration quotients of 1.40 mol O_2_/mol C, suggesting a lower H and O content, and higher O_2_ demand than commonly assumed. We found global variations in the O_2_ demand of organic matter respiration driven by population structure and cellular reorganisation under different environmental conditions. By testing how shifts in the macromolecular composition of phytoplankton cells affect the ocean’s O_2_ budget, we found that O_2_ consumption increases significantly when shifting cell composition from carbohydrate-rich to protein- or lipid-rich cells. As a result, low-O_2_ (hypoxic) zones in the ocean expanded by 75%. These findings demonstrate that cellular-level processes in marine phytoplankton shape the global O_2_ cycle and large-scale patterns of ocean biogeochemistry.

## Introduction

The ratios by which marine phytoplankton acquire inorganic elements and incorporate them into their cellular material (i.e., their elemental composition) effectively couples the global cycling of carbon (C), nitrogen (N), phosphorus (P), trace elements, and oxygen (O), and, therefore, play a fundamental role in shaping biogeochemical cycles. The relative ratio of N:P in seawater and phytoplankton distinguishes between nitrate and phosphate limitation, and regulates N_2_ fixation and denitrification rates^1–7^. The ratio of C to the limiting nutrient dictates the amount of atmospheric carbon dioxide (CO_2_) that can be sequestered in the deep ocean via the “biological pump”^8–10^. The hydrogen (H) and O content of the organic matter further provides an estimate of dissolve oxygen (O_2_) consumption and nitrate loss (denitrification) during aerobic and anaerobic respiration of organic matter^11,12^. Since organic matter respiration is the main route by which dissolved O_2_ is lost from the ocean interior^13,14^, understanding the composition of planktonic cells and the factors controlling it is crucial for understanding the O_2_ budget in the ocean.

Two metrics are often used to quantify O_2_ loss during organic matter respiration. The first is O_2_ demand, which expresses the amount of O_2_ consumed per unit P of organic matter (−ΣO_2_:P; mol O_2_/mol P). The second is the respiration quotient, which reflects the molar ratio of O_2_ consumed per CO_2_ released during organic matter respiration (−ΣO_2_:C; mol O_2_/mol P).

Traditionally, the stoichiometric equation representing the production and complete aerobic heterotrophic respiration of a phytoplankton cell (estimated from measured C:N:P of 106:16:1 in marine phytoplankton) is the Redfield–Ketchum–Richards (1963) (RKR) equation^15^:

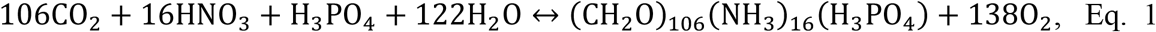

where production is the forward reaction and respiration is the reverse. Here, it is assumed that organic matter is composed mainly of carbohydrates (CH_2_O), ammonia (NH_3_), and phosphate (H_3_PO_4_). This assumption resulted in a “bulk” elemental composition of C_106_H_263_O_110_N_16_P, total O_2_ demand (−ΣO_2_:P) of 138 mol O_2_/mol P, and respiration quotients (−ΣO_2_:C) of 1.3 mol O_2_/mol C. However, phytoplankton are not made of only carbohydrates, but rather are protein-synthesising microorganisms, with protein accounting for 40**–**60% of phytoplankton cell weight^16^. Lipids and carbohydrates contribute 10 to 40% of cell weight (Figure *1*). Carbohydrates are a relatively oxidised form of organic matter, while proteins and lipids are more reduced (proteins contain reduced nitrogen, and both proteins and lipids contain CH_2_ chains). Therefore, the downstream biosynthesis of proteins and lipids from initially fixed carbohydrates releases O_2_, increasing the cell’s overall respiration quotient (Figure *1*b).

The O_2_ demand of organic matter can be estimated using different approaches. Using the average biochemical composition of the dominant macromolecules and their relative abundances in marine phytoplankton cells, Laws (1991) and Anderson (1995) have estimated an average O_2_ respiration demand of 150 mol O_2_/mol P and respiration quotient of 1.40 mol O_2_/mol C^17,18^. This “bottom-up approach” relies on culture-based laboratory estimates of phytoplankton composition and assumes a fixed average stoichiometry, without accounting for natural variability in species and environmental conditions. In contrast, “top-down approaches” rely on field measurements of dissolved nutrient ratios in the ocean. These measurements capture the community-level oxygen consumption of phytoplankton and associated organic matter (including particulate and labile dissolved pools); however, they are limited in time and space. Using such approaches, some studies have suggested that O_2_ demand is close to 140 mol O_2_/mol P^19,20^. However, other studies propose a higher O_2_ demand of ~170 mol O_2_/mol P^11,21,22^. A complication of this method is that it relies on assumptions about water mass transport and the initial concentrations of “preformed” nutrient and O_2_ concentrations (where “preformed” is defined as the concentration before subducting into the interior ocean). DeVries and Deutsch (2014) estimated water mass sources and their preformed values using a general circulation model constrained by biogeochemical tracers. They estimated that there are basin-scale variations in the −ΣO_2_:P, with higher values in the mid-latitude gyres and lower in equatorial and higher latitudes, and a global average of 147 mol O_2_/mol P^23^.

The global variations in the −ΣO_2_:P estimates of DeVries and Deutsch (2014) closely reflect variations in the C:P, N:P, and C:N ratios of particulate organic matter collected in the ocean^24^. Such variations are driven by variable environmental conditions such as light, temperature, and dissolved nutrient concentrations through changes in phytoplankton community structure and intracellular adjustments of macromolecular composition^24–27^.

While variations in C:P, N:P, and C:N of particulate organic matter are linked to −ΣO_2_:P, we currently lack an understanding of how the H and O content of organic matter shapes −ΣO_2_:P and −ΣO_2_:C at the global scale, and there are no systematic estimates of the global variations in the H and O content of particulate organic matter^28^. Nevertheless, we hypothesise that such variations exist and reflect differences in the underlying macromolecular composition of the living phytoplankton pool due to varying environmental conditions. For example, low-light conditions favour investment in light-harvesting proteins, which contain reduced N and CH_2_ chains, over C storage macromolecules such as carbohydrates and lipids (Figure *1*)^16,29^. Therefore, changes in environmental conditions may alter the complete elemental composition, including the H and the O content of freshly produced organic matter, and the amount of O_2_ consumed during respiration (i.e., the respiration quotients).

Here we address the following questions: (i) What is the average H and O content in marine phytoplankton, and how does this control O_2_ demand and the respiration quotient of organic matter? (ii) Are the H and O contents in marine phytoplankton fixed or vary spatially? If variable, what controls such variation? (iii) Similarly, are O_2_ demand and respiratory quotients constant, or do they vary spatially? What governs their variations? (iv) What is the sensitivity of the global O_2_ budget to variations in the H and the O content of marine phytoplankton?

To address these questions, we extended Laws’ (1991) and Anderson’s (1995) approach to the global ocean by using a macromolecular cellular allocation model embedded in the Darwin model^30,31^, a marine ecosystem model that simulates the dynamics of plankton and nutrient cycling within the Massachusetts Institute of Technology general circulation model (the MITgcm)^32^ framework. The cellular allocation model resolves variations in the major macromolecules of phytoplankton, such as proteins, carbohydrates, lipids, nucleic acids, and storage molecules (Figure *1*). These macromolecules vary as a function of environmental conditions^26,33^ (see Supplementary Information). By assuming predefined elemental ratios of each macromolecule (Figure *1*b), we resolved the elemental composition of phytoplankton biomass, including the H and the O content of phytoplankton cells, its O_2_ demand and respiration quotients. We also estimated the stoichiometry and O_2_ demand of particulate organic matter (POM), which is derived from phytoplankton and zooplankton biomass upon mortality, as well as from excess elements routed to POM due to selective grazing, under the assumption that grazers maintain elemental homeostasis (Supplementary Information).

## Results and Discussion

Overall, our findings reveal significant global variations in the H and the O content of phytoplankton cells and POM, as well as in their O_2_ demand and respiration quotient. By conducting sensitivity experiments, we found that the global ocean O_2_ budget is highly sensitive to variations in phytoplankton cells’ composition. These results provide a mechanistic understanding of the factors influencing the oxidation states of organic matter in the ocean and their impact on the global O_2_ cycle.

**Figure 1.**
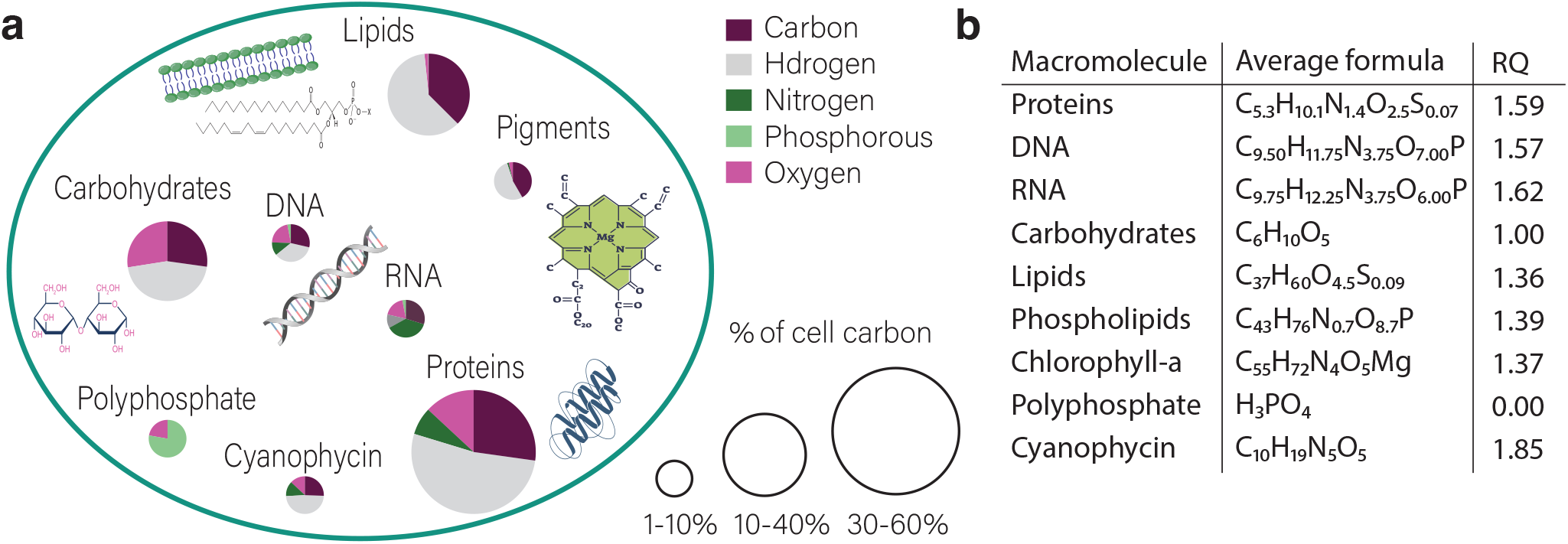
A schematic representation of a macromolecular composition of a phytoplankton cell. **(a)** Average composition of a phytoplankton cell. The size of the pie charts represents the estimated contribution of each macromolecular pool to cell C (mol C in macromol/mol C), while the proportions within the pie charts indicate the relative molar content of elements in each type of macromolecule. Adapted from Liefer et al., 2019^*51*^. **(b)** Average formula and respiration quotient (RQ) of major macromolecules identified in marine phytoplankton. The average formula and RQ of proteins, lipids, and nucleic acids are based on major amino acids identified in marine phytoplankton^*52–55*^, lipids identified in marine phytoplankton^*56,57*^, and an assumed equimolar distribution of the four deoxyribonucleotides and ribonucleotide bases, respectively (Extended Data Tables *1*—*2*).

### Average formula of a phytoplankton cell

We found an average phytoplankton cell composition emerges from the global simulations of C_107_H_190_N_16_O_53_P, with O_2_ demand of 149 mol O_2_/mol P, and a respiration quotient of 1.40 mol O_2_/mol C (Figure *2*a). This composition results from a modeled average composition of a phytoplankton cell consisting of 48% proteins, 23% lipids, 21% carbohydrates, 6% N-storage (as cyanophycin), 1.5% chlorophyll, and 0.4% nucleic acids (DNA + RNA) (mol C in macromolecules/mol C). Our results indicate a lower H and O content compared to the classical RKR formula. This is attributed to the inclusion of lipids and proteins in our simulations, which are more reduced than carbohydrates, which were not accounted for in the classical approach^15^.

We explored how much of the O_2_ consumption (149 mol O_2_/mol P) is due to (i) the oxidation of organic C as carbohydrates (CH_2_O); (ii) the oxidation of organic N to nitrate; and excess of H results from the inclusion of other macromolecules that are more reduced, mainly proteins and lipids (see Methods). Our model simulations suggest that 72% (107/149) of the O_2_ demand is due to the oxidation of C in the form of CH_2_O, 21% (32/149) is due to the oxidation of organic N to nitrate, and 7% (11/149) of the global O_2_ consumption is due to the oxygenation of excess of H due to the inclusion of variations in the lipid and proteins content of the organic matter (Figure *2*b). In comparison, the classical RKR equation assumes that 77% (106/138) of the O_2_ consumption is due to oxidising CH_2_O, and the remaining 23% (32/138) is due to the oxidation of organic N to nitrate.

**Figure 2.**
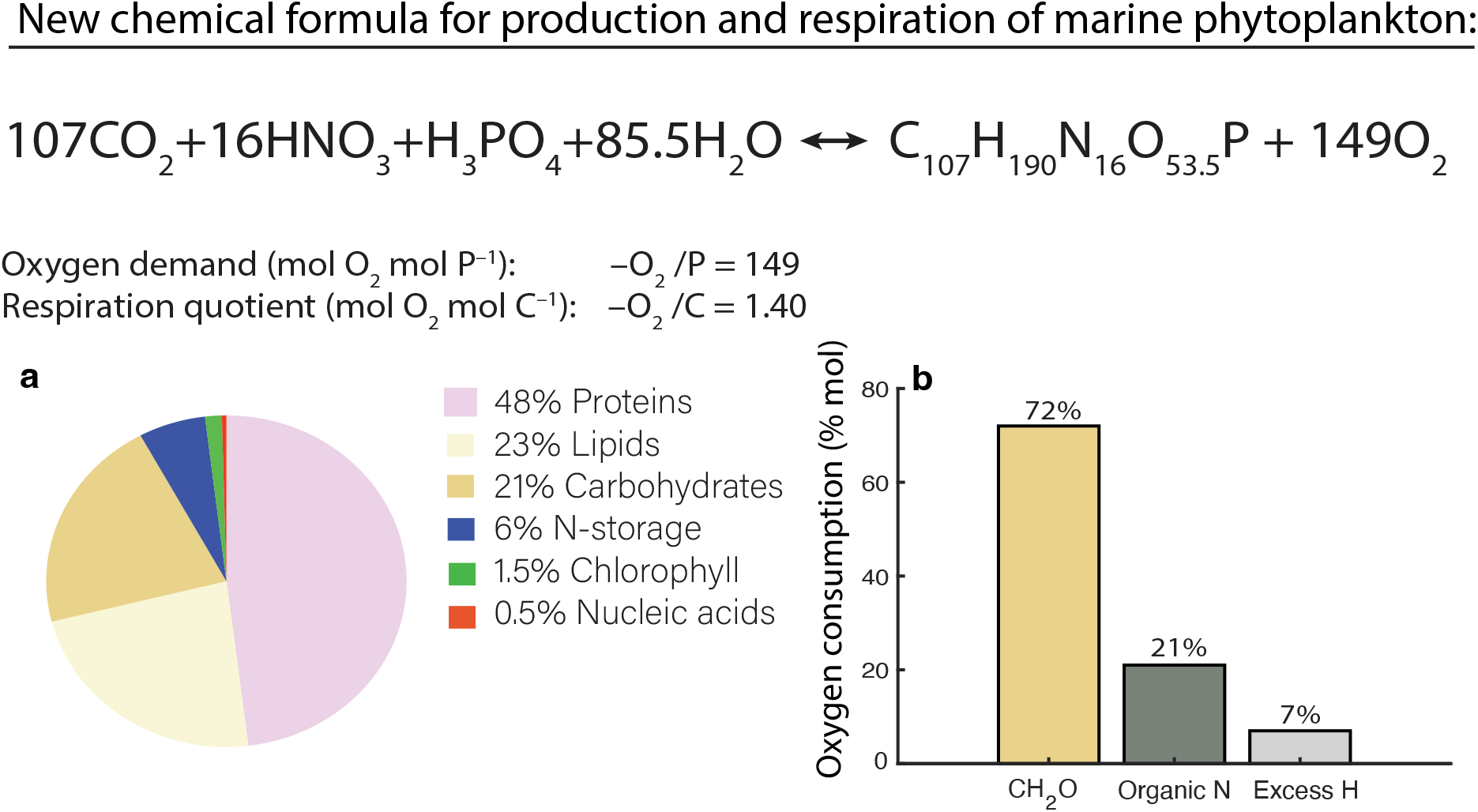
Average phytoplankton cell composition and associated oxygen demand. **(a)** Average chemical formula of production and heterotrophic respiration of marine phytoplankton derived from the model simulations, based on the macromolecular composition (mol C in macromol/mol C) presented in Panel **a. (b)** Relative contribution of each biochemical component to total O_2_ consumption during respiration (see Methods). For comparison, the classical Redfield-Ketchum-Richards (RKR) formula^*15*^ assumes the following formula: 106CO_2_ + 16HNO_3_ + H_3_PO_4_ + 122H_2_O ↔ (CH_2_O)_106_(NH_3_)_16_(H_3_PO_4_) + 138O_2_, with a total O_2_ demand of 138 mol O_2_/mol P and a respiration quotient of 1.30 mol O_2_/mol C, where 77% of the O_2_ consumption is due to oxidizing CH_2_O and 23% is due to the oxidation of organic N (see main text).

### Global variations in total particulate organic matter composition

Our model simulations further predict latitudinal trends in the elemental composition of total particulate organic matter in the surface ocean, encompassing both phytoplankton cells and detrital particulate organic matter (Figure *3*). Notably, we find that elemental ratios relative to P are primarily a result of phytoplankton community structure. Small phytoplankton have a lower storage capacity of phosphate relative to large phytoplankton^25,26,34^. Since phytoplankton in the ocean are limited mainly by light, iron, and nitrogen^35^, phytoplankton often store phosphate until saturation. Therefore, in nutrient-poor (oligotrophic) regions such as the subtropical gyres, where small phytoplankton with higher nutrient affinity and lower phosphate storage dominate, the C:P, H:P, N:P, and O:P ratios of total particulate organic matter are elevated (Figure *3*). On the other hand, in the subpolar regions and along the equator, where upwelling water supplies nutrients to the surface ocean, larger phytoplankton with higher maximum growth rates dominate. Larger phytoplankton have a higher phosphate storage capacity; therefore, the N:P, C:P, O:P, and H:P ratios of total particulate organic matter in these regions are lower.

Additionally, our model simulations suggest that the variations in N:C and H:C of total particulate organic matter reflect changes in cellular allocation to proteins, which have a significant amount of reduced nitrogen, resulting in relatively high N:C and H:C ratios of 0.26 and 1.90, respectively (Figure *3*). Therefore, in locations where cellular investment in proteins is high, N:C and H:C of total particulate organic matter is overall higher (Figure *3*c, i), which typically occurs in nutrient-rich, low-light conditions at high latitudes. There, N:C and H:C are higher as cells invest more in photosynthetic and biosynthetic proteins, which allow for more efficient light capture and faster growth^27,36^. Our modeled latitudinal trends in C:P, N:P and N:C of total particulate organic matter agree with available global observations (Figure *3*d–f).

**Figure 3.**
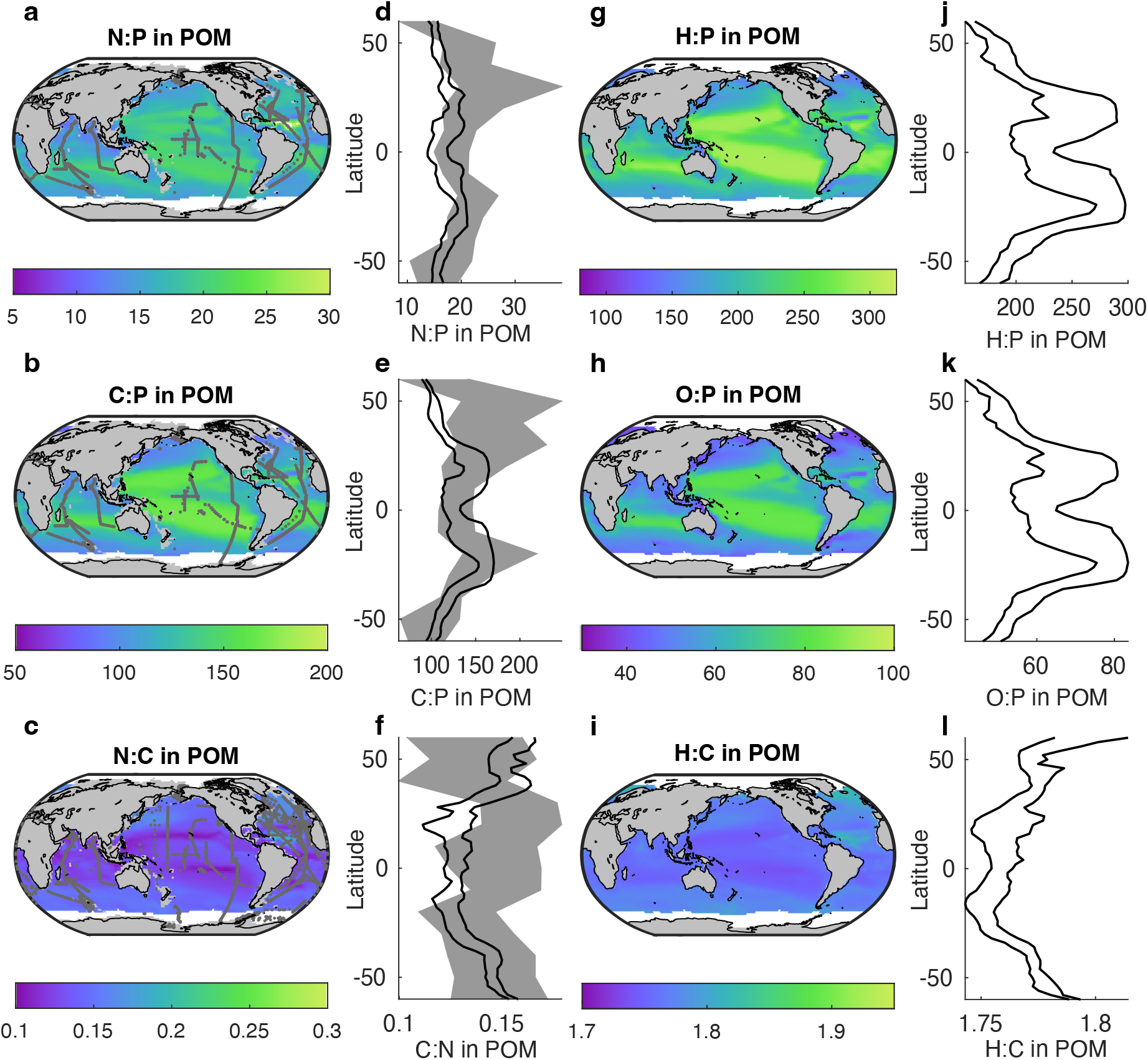
Global trends in the elemental composition of total particulate organic matter encompassing both phytoplankton and particulate organic matter. Maps show model-simulated, depth-integrated (0–170 m), annual mean molar ratios of **(a)** N:P, **(b)** C:P, **(c)** N:C, **(g)** H:P, **(h)** O:P, and **(j)** H:C in total particulate organic matter. Grey points in panels **a**–**c** mark observational sampling locations of particulate organic matter elemental composition from previous studies^*24,58*^. White areas in the global maps denote area with >50% annual sea ice cover. Panels **d–f** show latitudinal trends of observed (grey shading, 25–75th percentile range) and modeled (black lines, 25–75th percentile range) elemental ratios: **(d)** N:P (*n*_obs_=7,276), **(e)** C:P (*n*_obs_=6,983), and **(f)** N:C (*n*_obs_=43,639), respectively. Panels **j–l** show modeled latitudinal trends of **(j)** H:P, **(k)** O:P, and **(l)** H:C in total particulate organic matter, with black lines indicating the 25–75th percentile range of model values.

### Global variations in the oxygen demand and respiration quotients of total particulate organic matter

Our model simulations predict global variations in the O_2_ demand (−ΣO_2_:P) ranging from 110 to 280 mol O_2_/mol P, with higher values in the mid-latitude oligotrophic gyres and lower values at higher latitudes (Figure *4*a). Our results are within the uncertainty envelope of −ΣO_2_:P estimated using observations of dissolved nutrient ratios and a global circulation model (dashed-line Figure *4*c)^23^. Our model simulations further show that these variations primarily reflect changes in the C:P of total particulate organic matter driven by phytoplankton community structure (dark-red area in Figure *4*c). To a lesser extent, these variations in the O_2_ demand are influenced by reduced N demand (green area in Figure *4*c) with a smaller contribution to excess H due to the

inclusion of variations in the lipid and protein content of the organic matter (grey area in Figure *4*c, see Methods).

We also find that there are global variations in the respiration quotients of organic matter (−ΣO_2_:C) between 1.35 and 1.50, with a biomass-weighted global mean of 1.40 (Figure *4*b). The respiration quotients are lowest in the subtropical gyres. Here, phytoplankton invest a high proportion of their biomass in carbohydrates, which are relatively oxidised forms of organic matter. On the contrary, the respiration quotients are highest at high latitudes where phytoplankton are mainly composed of proteins, which are a more reduced form of organic matter and contain, in many cases, reduced nitrogen^27,36^.

Our results generally fall within the uncertainty range of available measurements of respiration quotients Figure *4*d), though the limited data on respiration quotients make a robust assessment challenging. Nevertheless, our model simulations qualitatively capture global patterns of macromolecular composition across the ocean^27,36^. Furthermore, our model simulations also suggest that the respiration quotient increases with depth, as phytoplankton invest more in photosynthetic proteins in response to diminishing light availability (Figure *4*e). These results agree with vertical profile measurements of respiration quotients from the Sargasso Sea^37^. This provides confidence in our predictions, although further sampling of respiration quotients across oceanic gradients, depths, and high latitudes is needed to refine the model parameterizations.

### The impact of variations in cellular composition on the global oxygen budget

There are many regions in the ocean where O_2_ is too low to sustain aerobic life. The extent of these hypoxic zones depends on a balance between O_2_ supply through ventilation pathways of the global ocean circulation and consumption by organic C respiration at depth^14^. These hypoxic regions are expanding, and numerous modeling studies suggest they will likely expand more due to human-induced climate change^38–41^. Along with changes related to reduced thermal solubility, ventilation, and primary productivity^42^, changes in the underlying macromolecular composition of phytoplankton may occur, which can either mute or exacerbate the impact of warming^43^. For example, under nitrate-poor conditions, cells may reduce their protein levels and increase investments in carbohydrates and/or lipids^16,36^, changing their respiration quotients. In this context, we ask: What would be the impact on the global ocean O_2_ budget if phytoplankton cells altered their composition to be predominantly carbohydrates with respiration quotient equal 1.3? Conversely, how would the O_2_ budget be affected if the cellular organic matter became more reduced with respiration quotient equal 1.5, for example, by containing more lipids and reduced proteins?

To diagnose the impact of varying phytoplankton macromolecular composition and the corresponding respiration quotients on the oceanic O_2_ budget, we conducted sensitivity experiments (see Methods). In these experiments, we varied respiration quotient according to the following values: in the first experiment, we used the variable respiration quotients estimated by this study (Figure *4*b). In the second experiment, we used a spatially homogeneous constant respiration quotient equal to 1.3. This scenario represents an end member where a cell becomes fully composed of carbohydrates (like the RKR assumption; Eq. 1). In the third experiment, we used spatially constant respiration quotients of 1.5. The respiration quotient in this experiment represents the other extreme end member, where the cell contains more lipids and/or proteins. We ran each of these model simulations for 2,000 years, allowing the deep ocean biogeochemistry to adjust and reach a quasi-steady state (Extended Data Figure *1*). After 2000 years, there remains a minor drift in the ocean O_2_ inventory (~0.01 mmol O_2_/m^3^ per decade in mean concentration, Extended Data Figure *1*a), presumably due to a small number of extremely long-lived modes of the global ocean circulation^44,45^. However, since the drift occurs in the same direction and at a similar rate across all simulations, and since we compare each simulation relative to the default (first) experiment with variable respiration quotient, we consider the impact of this drift on our results to be negligible.

**Figure 4.**
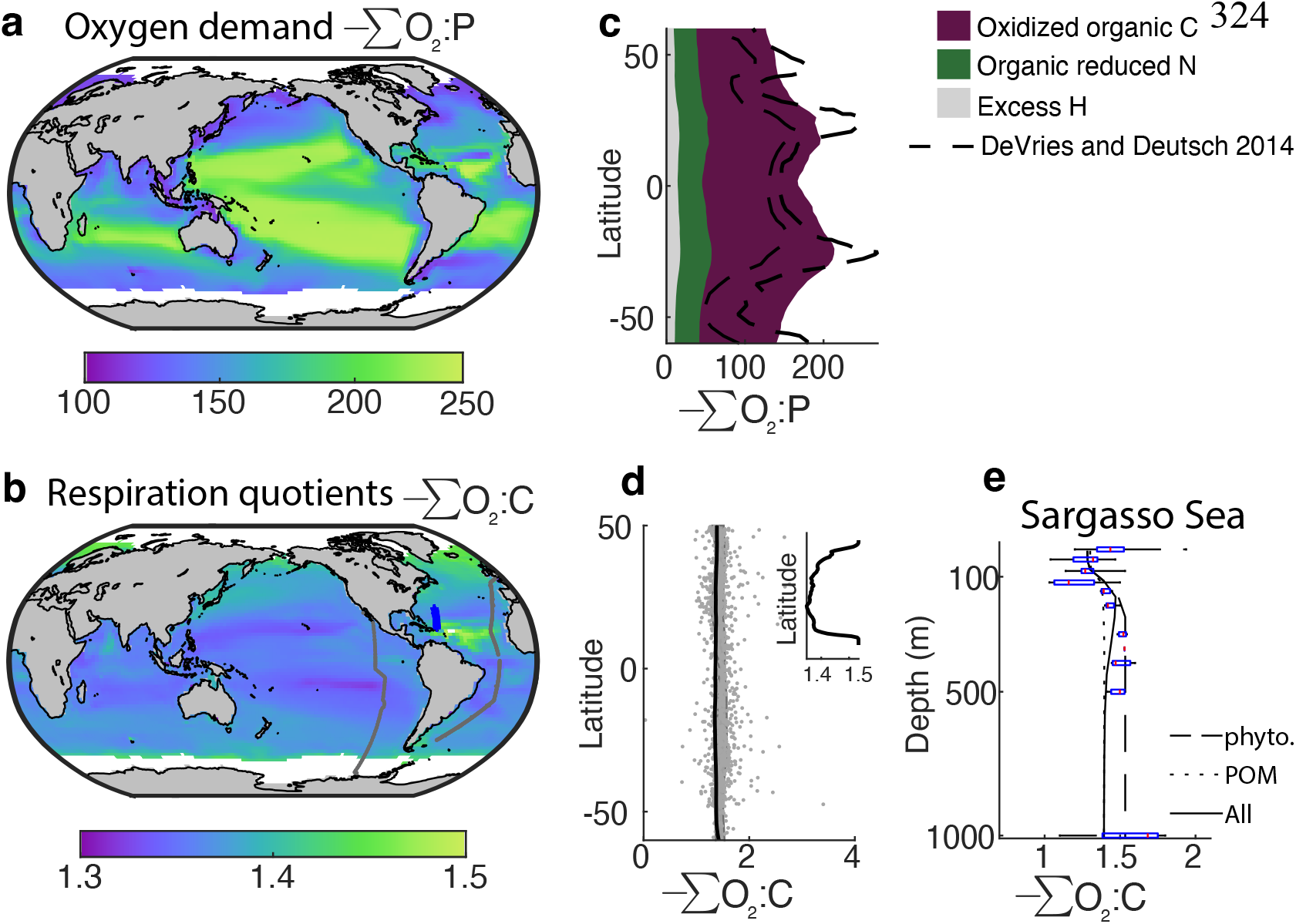
Oxygen demand and respiration quotients in total particulate organic matter encompassing phytoplankton and particulate organic matter. Model results of depth-integrated (0–170 m), annual means of **(a)** O_2_ demand (−ΣO_2_:P, mol O_2_/mol P) and **(b)** respiration quotients (−ΣO_2_:C, mol O_2_/mol C). The geographic locations of sampling stations in Panel **b** are marked as grey dots, with those analyzed in Panel **e** highlighted in blue. **(c)** The latitudinal trend in oxygen demand (−ΣO_2_:P) for the oxidation of organic C (dark-red area), organic reduced N (green area), and excess hydrogen (grey area) estimated by this study, along with estimated (dashed-black lines) values from DeVries and Deutsch (2014)^23^. **(d)** Latitudinal means of modeled (solid black lines, highlighted in the inset) and observed^*37,47,59*^ (grey dots) respiration quotients (n_obs_=1,028) of particulate organic matter. Observed respiration quotient includes O_2_ consumed during ammonium oxidation (PCOD+ 2PON)/POC). **(e)** Comparison between modeled and observed respiration quotient across depth in the Sargasso Sea. The modeled respiration quotient is represented by three components: the phytoplankton respiration quotient (dashed black line), the POM respiration quotient (dotted black line), and the biomass-weighted respiration quotient combining phytoplankton and POM (solid black line). Observations^*37*^ are grouped by depth, with box plots showing the 25th, 50th, and 75th percentiles and whiskers covering 99.3% of the data.

Our model simulations suggest that if cells globally become predominantly composed of carbohydrates with a respiration quotient of 1.3, global ocean inventory increases by ~5%, and the volume of the hypoxic and suboxic regions (defined here as regions where dissolved O_2_ concentrations fall below 62 and 5 μM, respectively^46^) decreases by 25% and 40%, respectively (compare orange and grey bars in Figure *5*a—c). Conversely, when cellular content becomes globally more reduced with a respiration quotient of 1.5 (for example, by containing more lipids or reduced proteins), the ocean holds 5.2% less O_2_, and the volume of hypoxic and suboxic regions increases by 30% and 115%, respectively (compare pink and grey bars in Figure *5*a—c).

Such a reduction in ocean O_2_ inventory by 5.2% when increasing respiration quotient by 0.1 is comparable to a projected reduction in global O_2_ inventory by the end of the 21^st^ century under the “business as usual” climate change scenario SSP-8.5^41,47^. Overall, shifting phytoplankton cellular composition from fully carbohydrate-based to fully protein-or lipid-based would result in a 75% increase of the volume of the hypoxic zones where [O_2_]<62 μM and more than a threefold increase in the suboxic zones where [O_2_]<5 μM (compare orange and pink bars in Figure *5*b, c). The sensitivity of the ocean volume to respiration quotient is most significant in the suboxic regions where [O_2_]<5 μM. At this threshold, the slope of the cumulative distribution function of O_2_ concentration is steepest, and a small shift leads to a significant change in suboxic volumes (Extended Data Figure *4*). Although not explicitly resolved in the model (Methods), such an increase in the extend of the suboxic regions will likely increase denitrification rates and the loss of nitrate through anaerobic respiration of organic matter.

**Figure 5.**
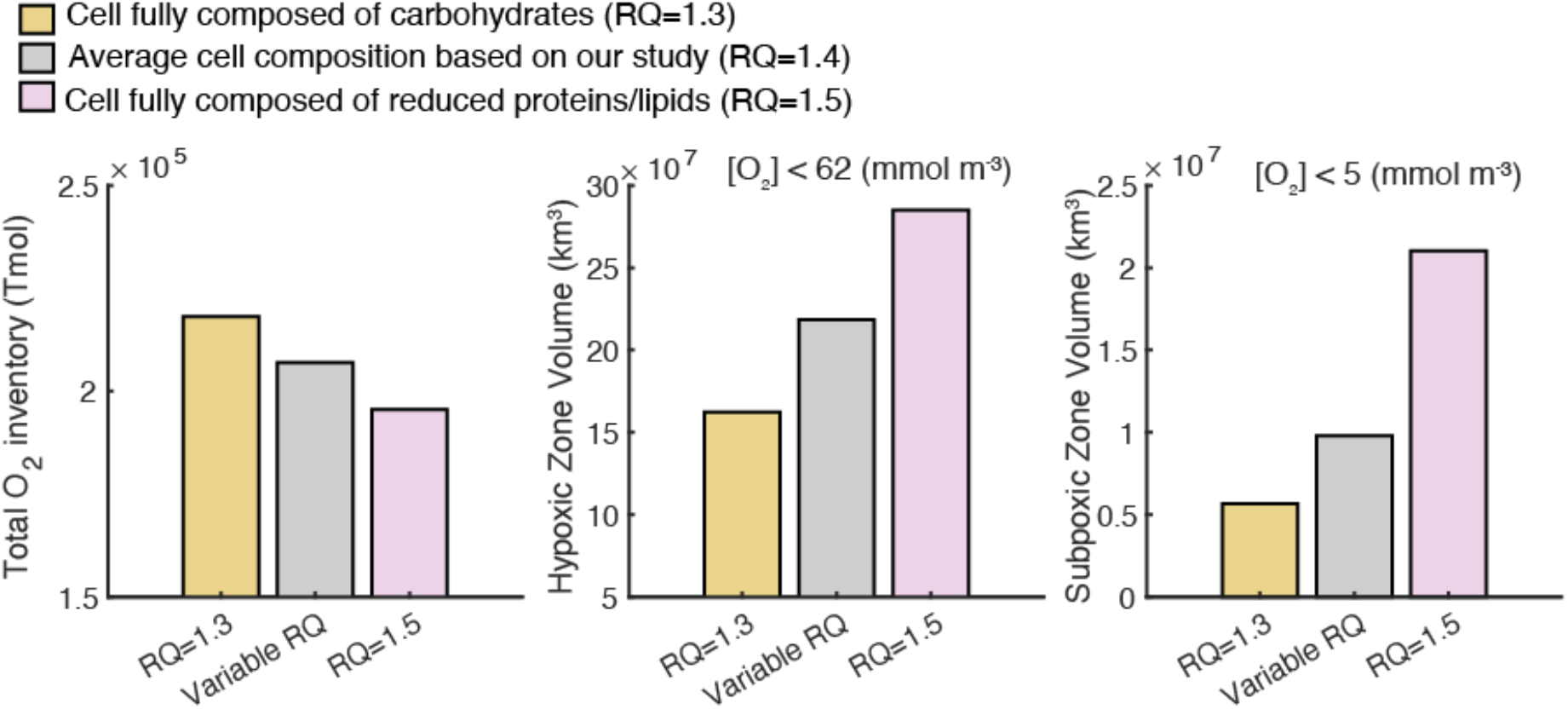
The sensitivity of the global oxygen budget and the extent of the oxygen hypoxic and suboxic zones to respiration quota. **(a)** Total O_2_ inventory (Tmol), **(b)** the volume of ocean hypoxic zones (km^3^) where [O_2_] < 62 μM, and **(c)** the volume of ocean suboxic zones (km^3^) where [O_2_] < 5 μM. Each bar chart comes from a different 2,000-year spin-up simulation with (i) the variable respiration quotients estimated by this study (grey), with (ii) respiration quotients of 1.3, assuming a cell is fully composed out of carbohydrates (orange), and (iii) respiration quotients of 1.5, assuming the cell containing more lipids and reduced proteins (pink).

## Conclusions

Our approach bridges the bottom-up and top-down methods for estimating the O_2_ demand of organic matter. By explicitly simulating variations in a phytoplankton cells’ macromolecular allocation under variable environmental conditions, we estimated the elemental composition, including H and O, O_2_ demand (−ΣO_2_:P), and respiration quotient (−ΣO_2_:C) of marine phytoplankton in the global ocean. We found an average chemical formula of production and heterotrophic respiration of marine phytoplankton cells of: C_107_H_190_N_16_O_53_P + 149O_2_→107CO_2_ + 16HNO_3_ + H_3_PO_4_ +85.5 H_2_O. This formula suggests that freshly produced phytoplankton cells contain less H and O and, therefore, are more reduced than the classical RKR equation, a formula often used in biogeochemical studies, which assumes that phytoplankton are predominantly composed of carbohydrates (Eq. 1). As a result, we estimated a higher O_2_ demand and respiration quotients of 149 mol O_2_/mol P and 1.40 mol O_2_/mol C, respectively. Importantly, we find that environmental variability gives rise to global variations in the elemental composition and O_2_ demand. Elemental ratios relative to P and −ΣO_2_:P are driven by variations in phytoplankton community structure, while elemental ratios relative to C and −ΣO_2_:C mostly resemble variations in the macromolecule composition. Our approach indicates that the ratio of dissolved O_2_ to regenerated nutrients is not fixed and can be predicted from first principles, providing a mechanistic link between phytoplankton biochemical properties (bottom-up) and large-scale biogeochemical patterns (top-down).

We further show that changes in cellular composition and the resulting shifts in cell respiration quotient significantly impact global O_2_ concentrations and the extent of the suboxic and hypoxic zones. Therefore, alterations in cellular composition under variable environmental conditions may either mitigate or amplify ocean deoxygenation. For example, the geologic record indicates that events of ocean anoxia were often linked to periods of abrupt warming, reduction in O_2_ solubility, and injection of nutrients from continental weathering that enhances export productivity and O_2_ consumption (e.g., the early Aptian and the Cenomanian-Turonian boundary which occurred ~120.5 and ~93.5 million years ago, respectively^48,49^). Under enhanced nutrient input, cells may favour proteins to facilitate growth. Higher content of reduced proteins can increase cells’ respiration quotients and amplify the extent of the O_2_ depletion zones.

On the other hand, the expansion of the mid-latitude oligotrophic gyres projected under anthropogenic warming^50^ may increase the relative fraction of carbon-storage macromolecules such as carbohydrates at the expense of proteins. This will shift the organic matter toward lower respiration quotients. This could partially mute the decline in oceanic O_2_ content expected with global warming, as less O_2_ will be consumed during respiration. Overall, our study highlights the importance of cellular-level processes in shaping large-scale patterns of ocean deoxygenation, which span hundreds of square kilometres and impact the metabolism of marine aerobic life.

## Methods

### Estimating oxygen demand from bulk elemental composition

To estimate oxygen demand from bulk elemental composition (C_a_H_b_O_c_N_d_P) we followed the approach of Paulmier et al., 2009^12^. Organic matter is traditionally represented by the following formula: (CH_2_O)_x_(NH_3_)_y_H_3_PO_4_. To account for deviations from this idealised formula, particularly the excess hydrogen content, the formula can be rewritten as C_x_(H_2_O)_w_H_z_(NH_3_)_y_H_3_PO_4_. The oxygen required to oxidise both the reduced carbon (x) and the excess hydrogen (z) can be calculated as follows:

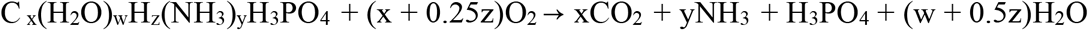

The elemental substitutions used are as follows:

- **x = a**, representing the number of carbon atoms,
- **w = c – 4**, assuming all oxygen not bound in H_3_PO_4_ is present as water (H_2_O),
- **y = d**, representing the number of nitrogen atoms (assumed to be in the form of NH_3_),
- **z = b – 2w – 3d – 3 = b – 2c – 3d + 5**, where the excess of hydrogen is calculated as the total hydrogen pool (b) minus the hydrogen in water (2w), ammonia (3d), and polyphosphate (3).

Ammonium oxidation proceeds through the following reactions:

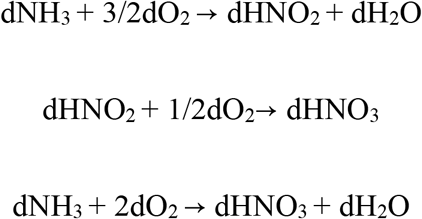

Thus, ammonium oxidation consumes 2d moles of O_2_.

The total oxygen demand for full oxidation of organic matter is given by:

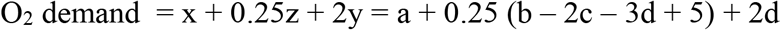

Which simplifies to:

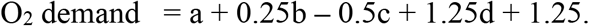

Here:

1. **a** is the amount of oxygen required to oxidise carbon (as CH_2_O),
2. **2d** accounts for oxygen required to oxidised reduced nitrogen,
3. **0.25b – 0.5c – 0.75d + 1.25** accounts for the oxygen required to oxidise excess hydrogen.

### The macromolecular model

The macromolecular model used here has been previously described^26^. The model resolves the growth rate and the cellular allocation of phytoplankton *j* to major macromolecules as a function of light and external nutrient concentrations. The major macromolecules resolved in the model are proteins, carbohydrates, lipids, nucleic acids, chlorophyll, and nutrient storage. See Supplementary Information for model development and equations.

### Ecosystem model

We simulate the biomass of two phytoplankton, representing small, prokaryotic-like phytoplankton (0.41 µm^3^ cell volume) with high nutrient affinity and low growth rate, large eukaryote-like phytoplankton (40 µm^3^ cell volume) with high growth rate and low nutrient affinity, and one grazer. The two phytoplankton types differ in their maximum photosynthesis rates, maximum nutrient uptake rates, nutrient half-saturation constants, sinking rates, and palatability to grazers (Supplementary Information). The equations governing phytoplankton and zooplankton carbon (*c*_*j*_), phosphate (*p*_*j*_), nitrogen (*n*_*j*_), and iron (*fe*_*j*_), all in mmol/m^3^ are:

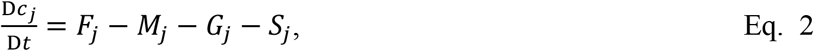

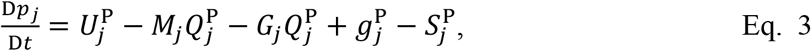

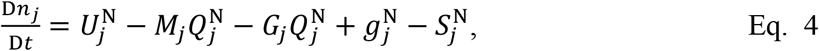

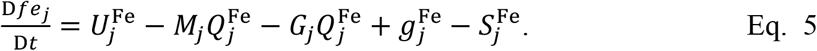

Here, D/Dt represents the material derivative that includes tracer transport, *F* represents phytoplankton net carbon fixation rates, which are calculated from the macromolecular model, *U* is nutrient uptake, *M* is the non-grazing mortality rate, *G* and *g* represent loss and gain from grazing, respectively, and *S* is sinking rates. Cellular quotas are defined as 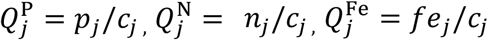 (mol/mol C). *F* = *U* = *G* = *S* = 0 for zooplankton, and *g* = 0 for phytoplankton. Symbol definitions and units are in Extended Data Table *3*.

### Dissolved nutrients

The concentrations of dissolved inorganic carbon, phosphorus, nitrogen, and iron (DIC, DIP, DIN, and DIFe, respectively) are governed by the following equations:

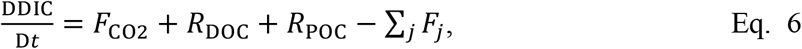

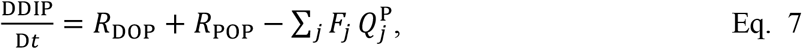

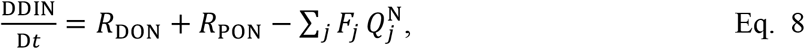

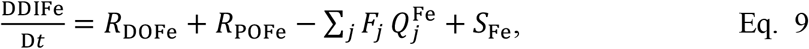

where *F*_CO2_ is the air-sea gas exchange, *R* is respiration rates, *F*_*j*_ is the net carbon fixation rates, and *S*_Fe_ is external iron sources (Supplementary Information).

### Particulate and dissolved organic matter

The model equations governing the formation, remineralisation, and sinking of particulate (PO*i*) and dissolved (DO*i*) organic matter (where *i*=C, H, N, O, P, Fe) are:

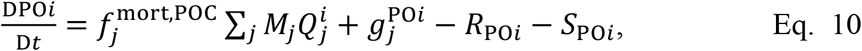

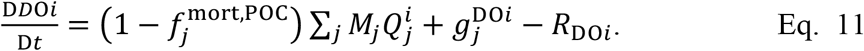

Here 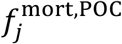 is the fraction of phytoplankton and zooplankton mortality that goes the particulate pool, *g* is gain from grazing, *R* is respiration rate, and *S* is sinking rate. Cellular quotas 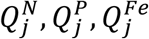 are computed from equations 2–5. 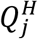 and 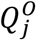 are estimated from the macromolecular model (see section below).

### Cellular oxygen and hydrogen

Cellular pools of organic hydrogen 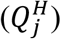 and oxygen 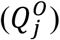 are estimated from the macromolecular model:

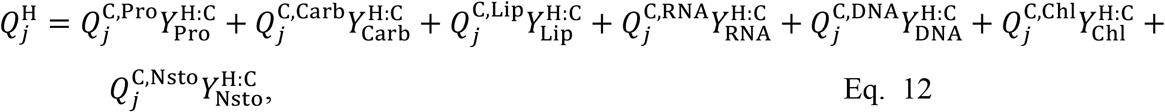

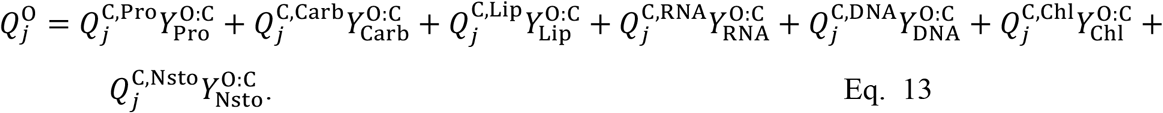

Here 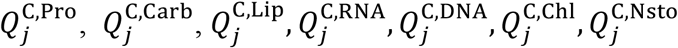 are cellular allocation to proteins, carbohydrates, lipids, RNA, DNA, chlorophyll, and N-storage molecules (mol C in macromolecule/mol C), respectively, calculated from the macromolecular model (Supplementary Information). 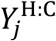 and 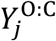 are the ratios of H:C and O:C in the different macromolecules (Figure *1*b).

### The oxygen cycle

The model equation governing the dissolved oxygen concentration is:

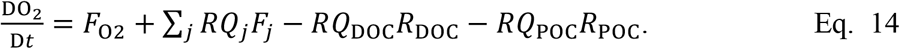

Here *F*_O2_ is the air-sea gas-exchange flux (mmol C m^−3^ day^−1^), which depends on gas solubility, wind speed, and temperature^60^, *F*_*j*_ is net carbon fixation rate (mmol C m^−3^ day^−1^), *R*_DOC_ and *R*_POC_ are respiration rates (mmol C m^−3^ day^−1^), and *RQ*_*j*_ (mmol O_2_ mmol C^-1^) is the respiration quotients of phytoplankton *j*:

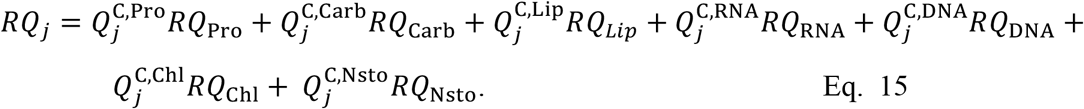

Where *RQ* is the respiration quotients of each macromolecule (Figure *1*b). The respiration quotients of the dissolved- and particulate-organic matter are estimated from their stoichiometry:

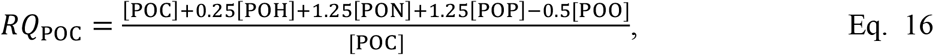

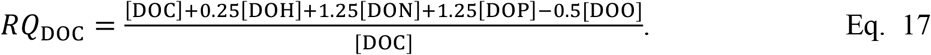

Here POC, POH, PON, POP, POO, DOC, DOH, DON, DOP, and DOO are the concentrations of dissolved and particulate organic carbon, hydrogen, nitrogen, phosphorus, and oxygen, respectively, calculated in equations 10—11.

### Physical model

The biogeochemical tracers are mixed and transported by the oceanic component of the MIT Integrated Global System Model (IGSM)^61–63^. The IGSM is an Earth system model of intermediate complexity, featuring a 2D (latitude, height) atmosphere and a 3D ocean circulation model (the MITgcm^32^) and is coupled to dynamic sea-ice and terrestrial models. The ocean component is configured here with a longitudinal resolution of 2.5° and a latitudinal resolution of 2°, and it has 22 non-uniform vertical depth levels, ranging from 10 m at the surface to 750 m at the bottom. We simulated the marine biogeochemical tracers for 2,000 years using “offline” circulation fields from the ocean component of the IGSM for preindustrial (1860) conditions. After this spinup, the biogeochemical tracers reach a quasi-steady state and are in reasonable agreement with global observations (Extended Data Figures *1*—*3*). The readers are referred to the citations above for further information.

### Sensitivity simulations

We conducted three sensitivity experiments to diagnose the impact of respiration quotients on the oceanic oxygen budget and the extent of the ocean minimum zones. In each of these simulations, the biogeochemical tracers were spun up for 2,000 years with the offline physical circulation fields described above, where all the parameters were kept constant, except the respiration quotients, which were set to the following values:

i. Spatially variable respiration quotients.
ii. Spatially constant respiration quotients of 1.3.
iii. Spatially constant respiration quotients of 1.5.

Case (i) is the spatially variable respiration quotient found by our dynamic model (Figure *4*b). Cases (ii) and (iii) represent two end-cases. In the second case, we assume that the cells are entirely composed of carbohydrates. This case is equal to the Redfield-Ketchum-Richards (RKR) assumption (Eq. 1), where the respiration quotient is spatially constant and equal to 1.3. In the third case, we assume a spatially homogeneous respiration quotient of 1.5. This scenario represents a situation where cells alter their composition to be predominantly composed of reduced lipids or proteins.

To calculate the global ocean inventory and the extent of the hypoxic and suboxic volumes (defined as regions where dissolved O_2_ concentrations are below 62 and 5 μM, respectively), we analysed the last 100 years of each simulation using MATLAB (Figure *5*).

### Uniqueness and limitations of the model

Our model does not include vertical migration of grazers, excretion of dissolved organic matter^64,65^, and accumulation of organic matter derived from bacteria^66^. Furthermore, we did not include preferential respiration of different macromolecules (e.g., some types of lipids greatly depleted with depth^67–69^). Follow-up work should consider such processes, as they can alter the depth at which oxygen is consumed, with implications for the global oxygen budget^37^.

We note that, our macromolecular model currently does not include N_2_ fixing phytoplankton. We therefore also did not include denitrification, where fixed nitrogen is lost to N_2_. Including denitrification without N_2_ fixation would result in a gradual depletion of nitrate over our long simulations. Our model simulations were designed to isolate the direct effect of respiration quotient on O_2_ inventory and provide a baseline from which these factors could be tested in future studies.

## Acknowledgments

SS would like to thank Dr. Miguel Frada and all members of the Darwin group for their fruitful discussions and comments. We thank the Interuniversity Institute for Marine Sciences in Eilat for hosting SS.

## Author contributions

Conceptualization: SS. Methodology: SS, KI, SD, OJ, MJF. Formal analysis SS. Investigation SS. Visualization: SS. Funding acquisition: SS, MJF. Data curation: SS, KI, SD, OJ, GLB. Supervision: MJF. Writing – original draft: SS. Writing – review & editing: MJF, KI, SD, OJ, GLB.

## Data and materials availability

All data needed to evaluate the conclusions in the paper are present in the paper.

## Data availability

The model outputs used in this study are available via Zenodo (DOI: 10.5281/zenodo.15719165 (I will only activate this link upon publication. For the review process, I can upload these netCDF files separately). The source code used for simulation and analysis is available official MITgcm GitHub repository. Custom modifications to the macromolecular allocation model are described in the Methods and Supplementary Information, and provided in the following link: https://github.com/shlomitsharoni/darwin3/tree/Darwin3_macromol.

## Extended Data Figures

**Extended Data Figure 1.**
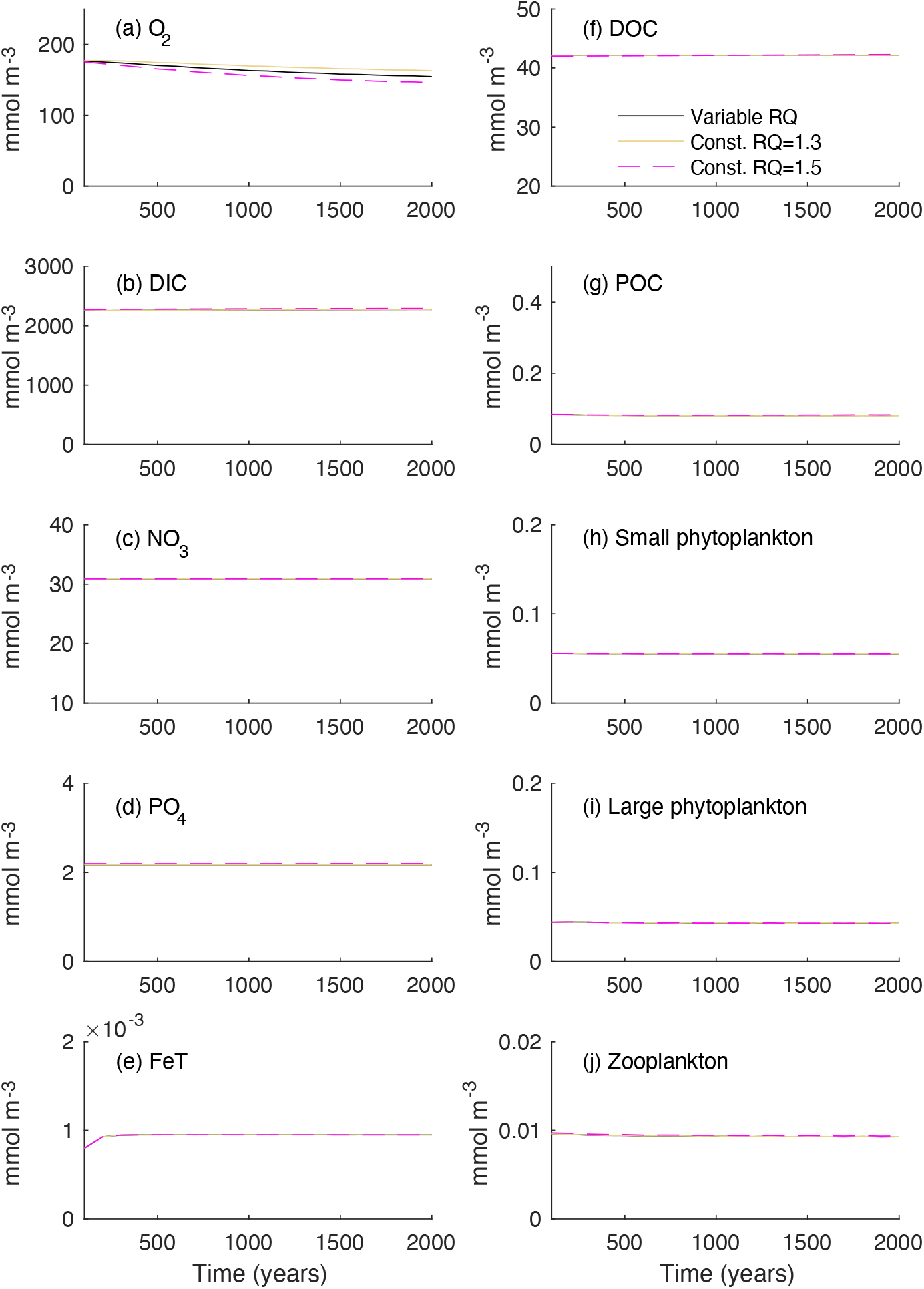
Modeled marine biogeochemical tracers. Concentrations (mmol/m^3^) of (**a**) dissolved oxygen (O_2_), (**b**) dissolved inorganic carbon (DIC), (**c**) dissolved nitrate (NO_3_), (**d**) dissolved phosphate (PO_4_), (e) total dissolved inorganic iron (FeT), (**f**) dissolved organic carbon (DOC), (**g**) particulate organic carbon (POC), (**h**) small phytoplankton organic carbon, (**i**) large phytoplankton organic carbon, and (**j**) zooplankton organic carbon. Solid black lines represent simulations with variable respiration quotients. A value of 45 mmol/m^3^ was added to the model DOC pool to represent non-reactive DOC. Black, pink, and orange lines represent simulations with the variable respiration quotients estimated by this study, with constant respiration quotients of 1.3 and 1.5, respectively.

**Extended Data Figure 2.**
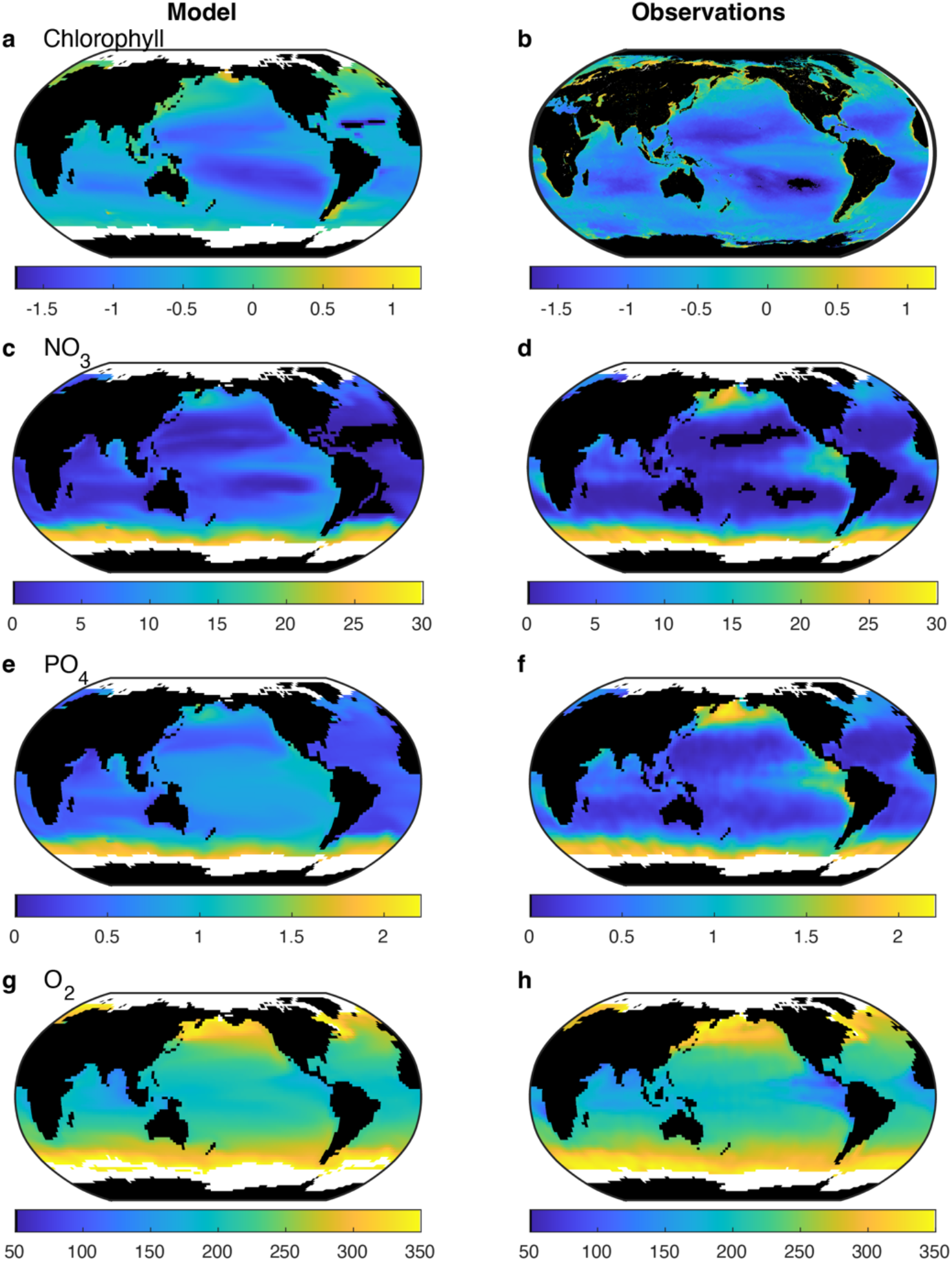
Comparison between observed and modeled biogeochemical tracers. Depth-integrated (0**–**115 m), annual means of (**a, b**) chlorophyll concentration (mg Chl/m^3^) (**c, d**) nitrate, (**e, f**) phosphate, and (**g, h**) oxygen. All tracers, except chlorophyll, are reported in mmol/m^3^. Observed chlorophyll data are derived from ocean colour products^*70*^, while dissolved nutrient observations are from the World Ocean Atlas^*71*^. The left column shows model results, and the right column shows observational data. The model results represent the 100-year mean of the model output for the year 2000 of simulation with variable respiration quotients.

**Extended Data Figure 3.**
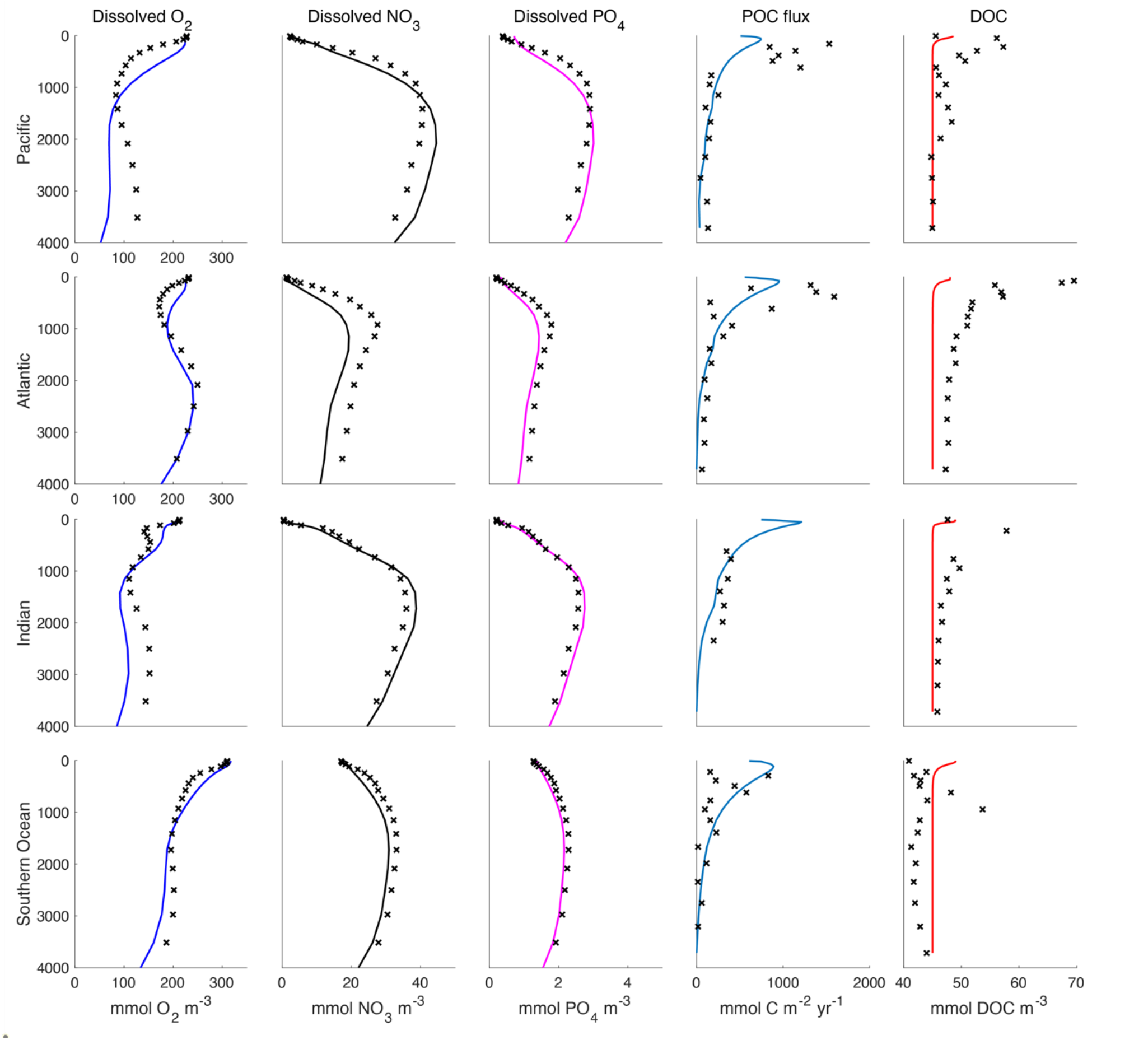
Comparison between vertical profiles of modeled and observed biogeochemical tracers. The comparison includes modeled (solid lines) and observed (black x’s) concentrations of dissolved oxygen, nitrate, and phosphate^*71,72*^, flux of particulate organic matter (POC; mmol C/m^2^/yr)^*73*^, and dissolved organic matter (DOC) concentration^*74,75*^ across different ocean basins. All tracers, except POC flux, are reported in mmol/m^3^. The model results represent the 100-year mean of the model output for the year 2000 of simulation with variable respiration quotients. A value of 45 mmol/m^3^ was added to the model DOC pool to represent non-reactive DOC.

**Extended Data Figure 4.**
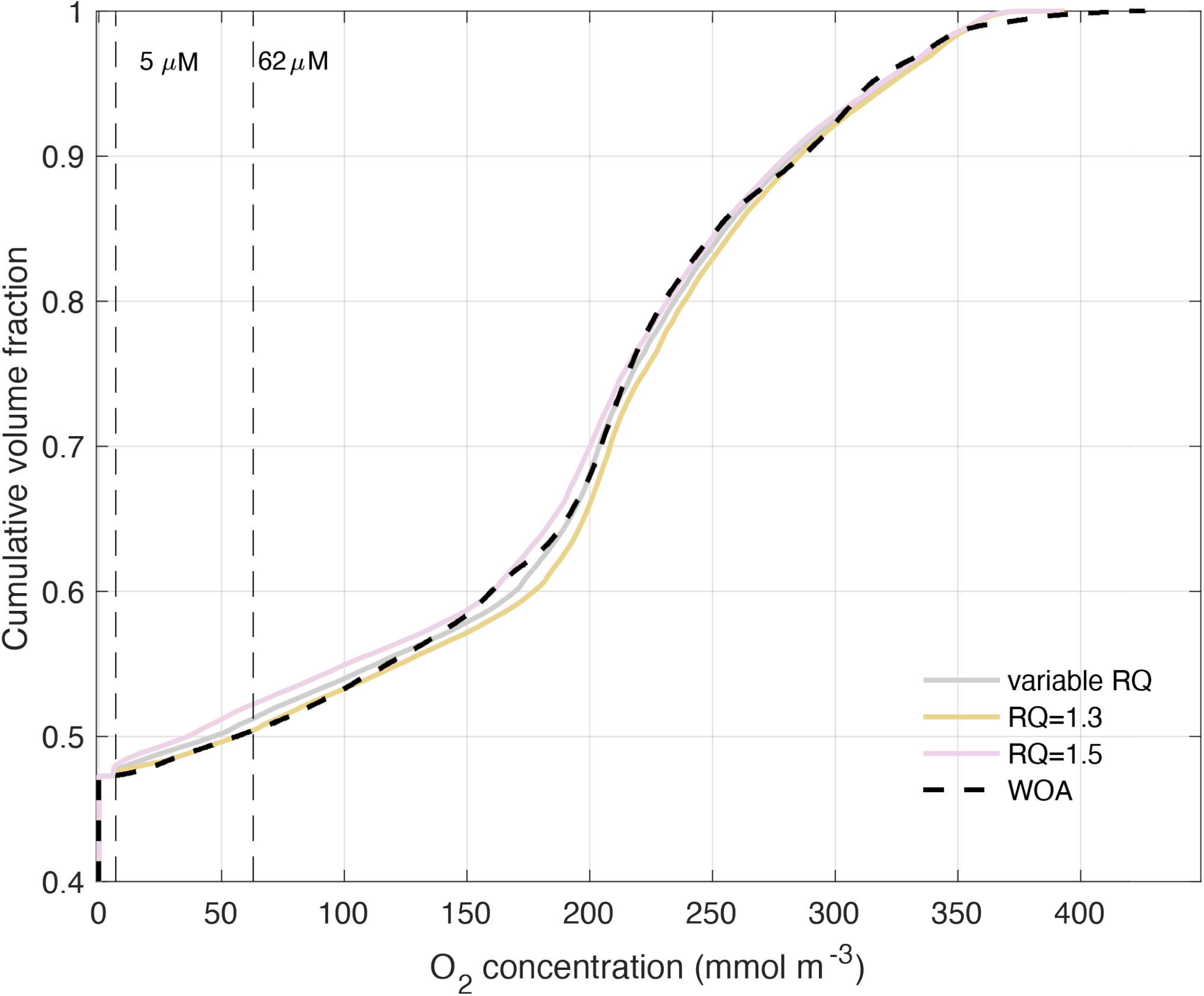
Cumulative distribution of ocean oxygen concentrations under different scenarios. The plot shows the cumulative volume fraction of ocean water as a function of O_2_ concentration (mmol/m^3^). Each line comes from a different 2,000-year simulation with (*i*) the variable respiration quotients estimated by this study (grey), with (*ii*) respiration quotients of 1.3, assuming a cell is fully composed out of carbohydrates (orange), and (*iii*) respiration quotients of 1.5, assuming the cell containing more lipids and reduced proteins (pink). Observational data from the World Ocean Atlas^*71*^ is shown for comparison. Vertical dashed lines mark the hypoxic and suboxic thresholds at 62 µM and 5 µM O_2_, respectively. The steepest portions of the curves indicate where small changes in oxygen leads to large changes in suboxic zone volume.

## Extended Data Tables

**Extended Data Table 1.**
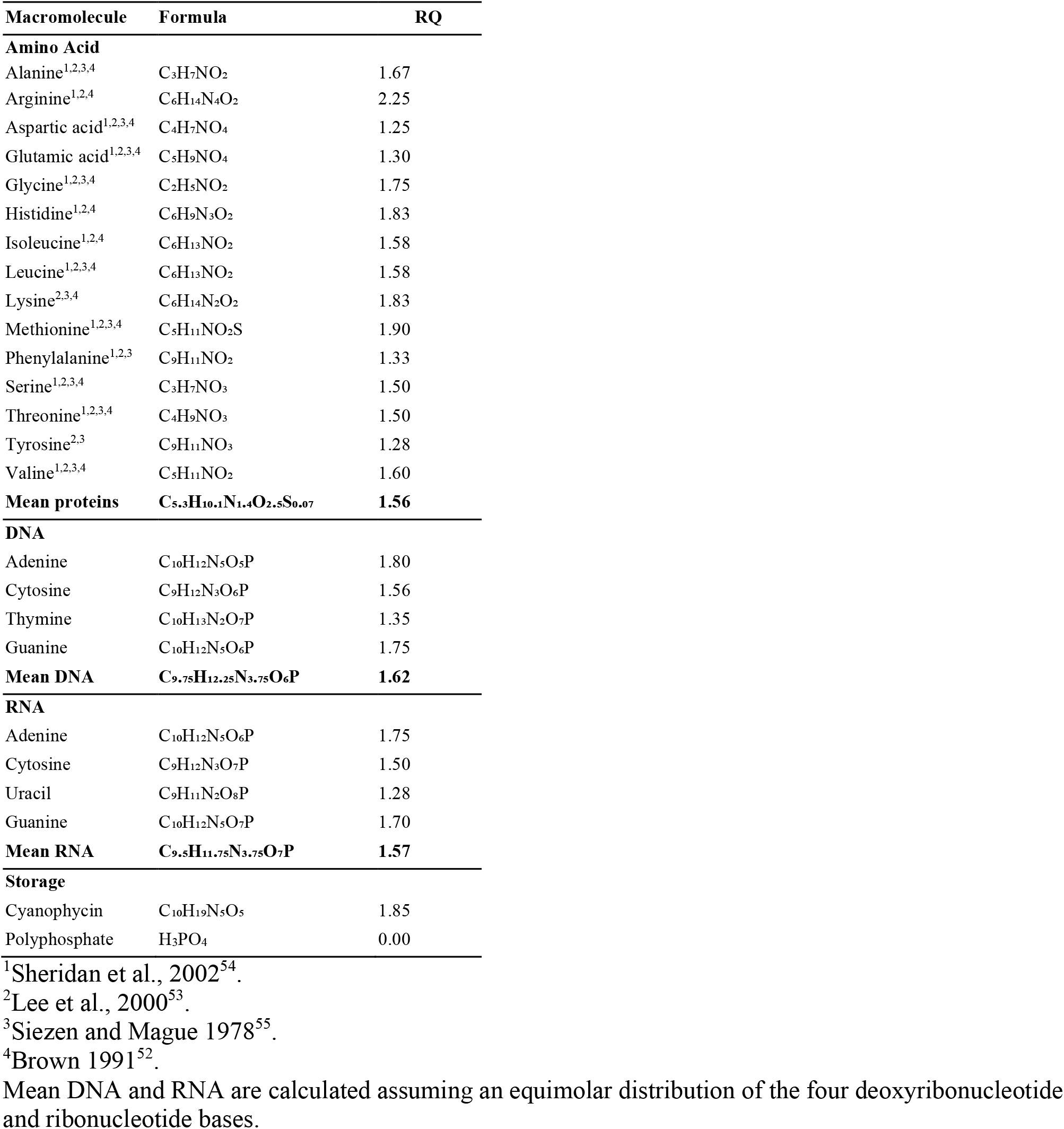
Elemental composition of amino and nucleic acids identified in marine phytoplankton.

**Extended Data Table 2.** Lipid types and their chemical formula.

| Lipids | Formula | RQ |
| --- | --- | --- |
| **Betaine lipid<sup>1</sup>** |  |  |
| Diacylglyceryl trimethylhomoserine (DGTS) | C<sub>46</sub>H<sub>81</sub>NO<sub>7</sub> | 1.40 |
| Diacylglyceryl hydroxymethyl-trimethyl-b-alanine (DGTA) | C<sub>49</sub>H<sub>84</sub>NO<sub>7</sub> | 1.38 |
| Diacylglyceryl carboxyhydroxymethylcholine (DGCC) | C<sub>51</sub>H<sub>84</sub>NO<sub>8</sub> | 1.35 |
| **Glycolipid<sup>1</sup>** |  |  |
| Monogalactosyldiacylglycerol (MGDG) | C<sub>44</sub>H<sub>74</sub>O<sub>10</sub> | 1.31 |
| Digalactosyldiacylglycerol (DGDG) | C<sub>49</sub>H<sub>85</sub>O<sub>15</sub> | 1.28 |
| Sulfoquinovosyldiacylglycerol (SQDG) | C<sub>41</sub>H<sub>73</sub>O<sub>12</sub>S | 1.34 |
| **Sterols<sup>2</sup>** |  |  |
| Fungisterol | C<sub>28</sub>H<sub>48</sub>O | 1.41 |
| Chondrillasterol | C<sub>29</sub>H<sub>48</sub>O | 1.40 |
| Brassicasterol | C<sub>28</sub>H<sub>46</sub>O | 1.39 |
| Chalinasterol | C<sub>28</sub>H<sub>46</sub>O | 1.39 |
| Desmosterol | C<sub>27</sub>H<sub>44</sub>O | 1.39 |
| **Quinones<sup>3</sup>** |  |  |
| Ubiquinone | C<sub>49</sub>H<sub>74</sub>O<sub>4</sub> | 1.34 |
| Plastoquinones | C<sub>53</sub>H<sub>80</sub>O<sub>2</sub> | 1.36 |
| Phylloquinol | C<sub>31</sub>H<sub>48</sub>O<sub>2</sub> | 1.35 |
| **Mean lipids** | **C<sub>37</sub>H<sub>60</sub>O<sub>4.5</sub>** | **1.36** |

<sup>1</sup>Van Mooy and Fredricks, 2010<sup>57</sup>.
<sup>2</sup>Piepho et al., 2012<sup>76</sup>.
<sup>3</sup>Becker et al., 2018<sup>56</sup>.

**Extended Data Table 3.**
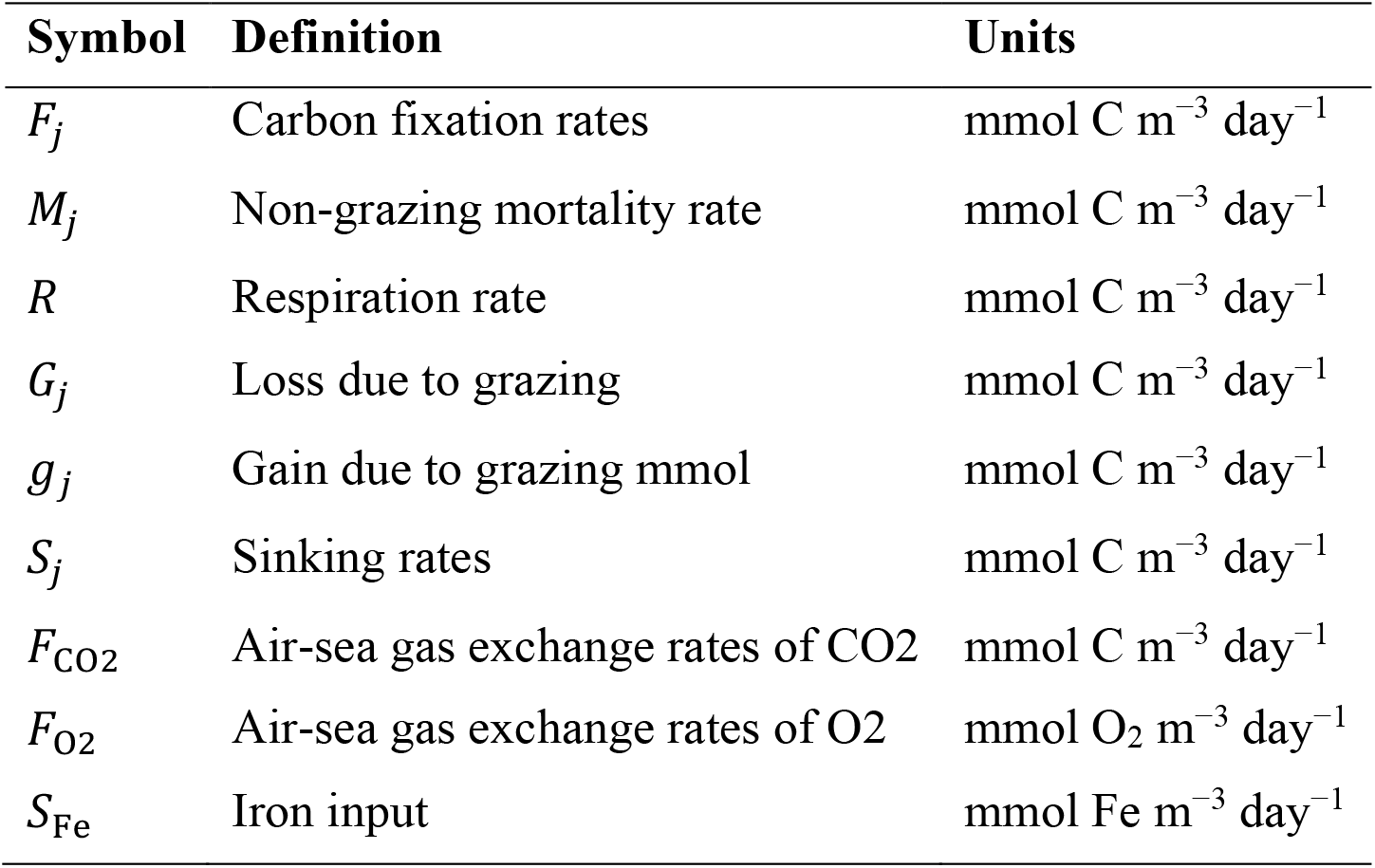
Model fluxes.

| Symbol | Definition | Units |
| --- | --- | --- |
| $F_j$ | Carbon fixation rates | mmol C m<sup>-3</sup> day<sup>-1</sup> |
| $M_j$ | Non-grazing mortality rate | mmol C m<sup>-3</sup> day<sup>-1</sup> |
| $R$ | Respiration rate | mmol C m<sup>-3</sup> day<sup>-1</sup> |
| $G_j$ | Loss due to grazing | mmol C m<sup>-3</sup> day<sup>-1</sup> |
| $g_j$ | Gain due to grazing mmol | mmol C m<sup>-3</sup> day<sup>-1</sup> |
| $S_j$ | Sinking rates | mmol C m<sup>-3</sup> day<sup>-1</sup> |
| $F_{CO2}$ | Air-sea gas exchange rates of CO2 | mmol C m<sup>-3</sup> day<sup>-1</sup> |
| $F_{O2}$ | Air-sea gas exchange rates of O2 | mmol O<sub>2</sub> m<sup>-3</sup> day<sup>-1</sup> |
| $S_{Fe}$ | Iron input | mmol Fe m<sup>-3</sup> day<sup>-1</sup> |

